# Impaired early visual categorization of fear in social anxiety

**DOI:** 10.1101/702498

**Authors:** Melissa Meynadasy, Kevin Clancy, Jessica Simon, Wei Wu, Wen Li

## Abstract

Social anxiety is associated with biased social perception, especially of ambiguous cues. While aberrations in high-level processes, including cognitive appraisal and interpretation of social signals, have been implicated in such biases, contributions of early, low-level stimulus processing remain unclear. Categorical perception is known to be an efficient process to resolve signal ambiguity, and categorical emotion perception can swiftly classify sensory input, “tagging” biologically important stimuli at early stages of processing to facilitate ecological response. However, early threat categorization could be disrupted by exaggerated threat processing in anxiety, resulting in biased perception of ambiguous signals. We tested this hypothesis among individuals with low and high trait social anxiety (LSA and HSA), who performed a 2-alternative-forced-choice (fear or neutral) task on facial expressions parametrically varied along a neutral-fear continuum. Clear divergence between the groups emerged in the profiles of reaction time (RT) and early visual response along the neutral-fear continuum. The LSA group exhibited a RT profile characteristic of categorical perception with drastically increased RT from neutral to intermediate (boundary) fear intensities, contrasting monotonous, non-significant RT changes in the HSA group. Neurometric analysis along the continuum identified an early fear-neutral categorization operation (arising in the P1, an early visual event-related potential/ERP at 100 ms) in the LSA (but not HSA) group. Absent group differences in higher-level cognitive operations (identified by later ERPs), current findings highlight a dispositional cognitive vulnerability in early visual categorization of social threat, which could precipitate further cognitive aberrations and, eventually, the onset of social anxiety disorder.

---

The ability to detect danger and initiate defensive response is critical for the survival and well-being of an individual. In anxiety, however, these processes become maladaptive, characterized by exaggerated threat processing and responding (Gray & McNaughton, 2000; Lang, Davis, & Ohman, 2000). Such maladaptive responses are especially prominent in the face of ambiguous threat, causing various cognitive and affective impairments in anxious individuals (Clark & Wells, 1995; Eysenck, Mogg, May, Richards, & Mathews, 1991; Rapee & Heimberg, 1997; Richards & French, 1992). The social environment is particularly rich with subtle and ambiguous cues, and biases to such social signals represent a key aspect of the psychopathology of social anxiety (Amir, Foa, & Coles, 1998; Clark & Wells, 1995; Forscher & Li, 2012; W. Li, Zinbarg, Boehm, & Paller, 2008; Philippot & Douilliez, 2005; Rapee & Heimberg, 1997; Yoon & Zinbarg, 2007).

It is generally understood that threat processing involves multiple processes and stages or “waves” (Adolphs, 2002; LeDoux, 1995; Pessoa & Adolphs, 2010; Vuilleumier & Pourtois, 2007). Cognitive theories of anxiety have implicated aberrations in early and late stages of threat processing (Bar-Haim, Lamy, Pergamin, Bakermans-Kranenburg, & van, 2007; Beck & Clark, 1997; LeDoux, 1995; W. Li, 2019; Mogg & Bradley, 1998; Ohman, 1993). Much empirical evidence has been garnered with respect to later threat processing in social anxiety, such as biased appraisal and interpretation (Heinrichs & Hofmann, 2001; Mathews & MacLeod, 2005). However, largely elusive to behavioral observation, early pre-attentive processing of threat is less understood. Therefore, new insights into these early processes could shed important light on cognitive vulnerability to ambiguous cues in social anxiety.

Among early processes, stimulus categorization could be particularly relevant to the threat bias in social anxiety. Categorical perception is a powerful process to resolve stimulus ambiguity, by partitioning the sensory space with sharp boundaries to generate distinct object categories (Harnad, 1987). For example, in color perception, categorical perception dissects the continuous light spectrum with abrupt boundaries to generate discrete color percepts (e.g., green, blue; (Bornstein, Kessen, & Weiskopf, 1976). Importantly, categorical object perception occurs automatically and at a credible speed (i.e., commensurate with object detection; (Green & Fei-Fei, 2014; Grill-Spector & Kanwisher, 2005). Consistent with that notion, visual event-related potentials (ERPs), particularly the P1 component, have specified the latency of object categorization as early as ~100 ms (Thorpe, 2009; Thorpe, Fize, & Marlot, 1996).

In terms of emotion, categorical perception also assigns boundaries along continua of varying emotional expressions, classifying complex and ambiguous emotional stimuli into distinct basic types (e.g. fear, happy, neutral; (Calder, Young, Perrett, Etcoff, & Rowland, 1996; Etcoff & Magee, 1992). In addition, categorical emotion perception is known to occur automatically and swiftly as well (Campanella, Quinet, Bruyer, Crommelinck, & Guerit, 2002; Roberson & Davidoff, 2000), serving a critical ecological function by efficiently “tagging” a potential threat cue to prioritize threat analysis (Yantis & Johnson, 1990). Indeed, like object categorization, our recent neurometric analysis of ERP responses along a neutral-fear continuum has identified threat categorization in the P1 component at 100 ms (Forscher, Zheng, Ke, Folstein, & Li, 2016).

While categorical emotion perception can be highly advantageous, it could be impaired due to maladaptive threat processing in anxiety. That is, by exaggerating threat processing to the extent that even mild, dismissible signals are encoded as threatening (Mogg & Bradley, 1998), anxiety could blur the boundary between threat and non-threat, thus disrupting categorical perception of threat. As a result, compromised resolution of ambiguous cues could not only cause significant social impairment but also subject anxious individuals to a great deal of uncertainty and stress, thereby fueling and perpetuating anxiety symptoms. In this study, we set out to test the hypothesis of impaired threat categorization in social anxiety.

To uncover cognitive processes underlying emotion processing, many creative paradigms (e.g., emotional Stroop, dot-probe, and visual search) have been used. However, behavioral measures from these tasks are likely confounded by operations from multiple stages (McNally, 1995), and early processes (that are brief and remote from final behavioral output) are especially difficult to specify behaviorally. Electrophysiological research has the superb temporal resolution to dissociate stages of information processing but is nonetheless limited in ascribing specific cognitive functions to them. To tackle these challenges, we recently developed a method by combining psychophysical testing and neurometric modeling of ERP responses to parametrically manipulated fearful expressions (Figure 1A-B; (Forscher et al., 2016). As such, we identified four key cognitive operations in fear perception that unfold in sequence and map onto four ERP components—fear categorization (P1), detection (P300), valuation (early LPP), and conscious awareness (late LPP; Figure 1C). Here, deriving ERP metrics for these operations, we compared individuals with high and low trait social anxiety on behavioral and neural measures of fear perception, with a particular focus on categorical perception of fear.

**Figure 1:**
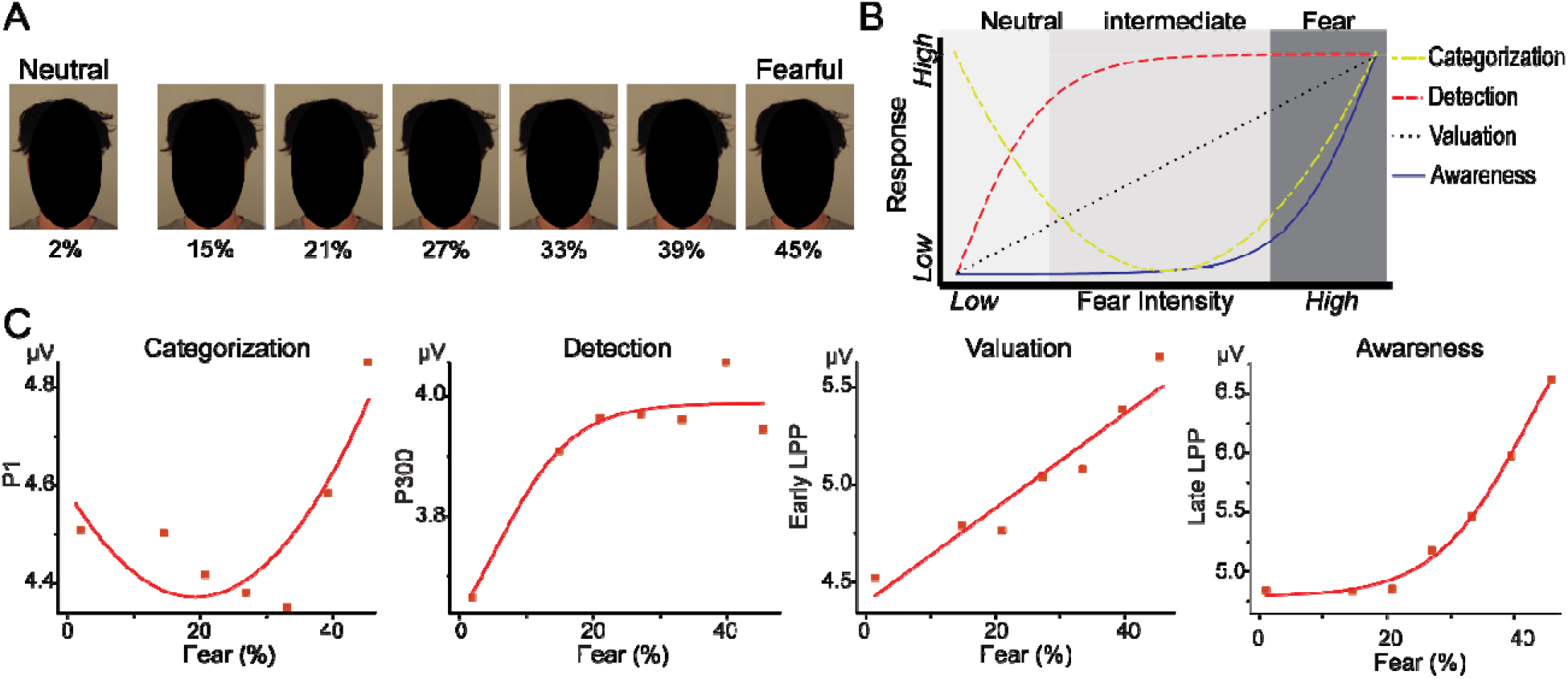
Experimental stimuli and four key processes of fear perception. (A) Example stimuli along the neutral-fear morph continuum with neutral (2%) and fear (45%) prototype levels and five levels in between (15%-39% fear in 6% increments). ***Note***: Per bioRxiv policy, faces are occluded for the current submission. (B) Psychological processes of interest are modeled according to their characteristic response functions: fear-neutral categorization by a quadratic function (yellow line); fear detection by a sigmoidal function (with an upper asymptote; red line); fear valuation by a linear function (gray line); and fear awareness by a sigmoidal function (with a lower asymptote, blue line). (C) Neurometric curve fitting identifies four key processes emerging over time in sequence. Fear-neutral categorization: P1 (at 100 ms) amplitudes conform to an upward quadratic function. Fear detection: P300 (at 300 ms) amplitudes conform to (the upper half of) a sigmoid function. Fear valuation: early LPP (400-500 ms) amplitudes conform to a linear function. Fear conscious awareness: late LPP (500-600 ms) amplitudes conform to (the lower half of) a sigmoid function. Adapted from Forscher, et al., 2016.

## Methods

### Participants

The data analyzed here belong to a larger study, which was initially reported in (Forscher et al., 2016). Forty-five undergraduate students (25 female; mean age 20 ± 4 years) participated in the study, with three excluded from ERP analyses due to excessive EEG interference and artifacts. All students were right-handed, with normal or corrected-to-normal vision, and had no history of neuropsychological problems and no current use of psychotropic medications. Further details about the participants (and experimental procedures) are presented in (Forscher et al., 2016).

#### Social Phobia Scale (SPS)

At the beginning of the experiment, we administered the SPS to measure trait social anxiety. The SPS is a commonly used self-report measure of sensitivity to social threat (Mattick & Clarke, 1998). It consists of 20 items pertinent to social situations (e.g., I am worried people will think my behavior odd; I become anxious if I have to write in front of other people), to which participants rated their *general* (trait) patterns with a 5-point scale ranging from 0 (very little) to 4 (very much). The high internal consistency (alpha coefficient = .95) for this sample indicated very strong reliability of the scale. Applying a median split of the SPS scores, we divided the sample into a high social anxiety (HSA) group and a low social anxiety (LSA) group. SPS scores were significantly higher in the HSA group [mean (SD) = 30.91 (13.34); range: 14 - 61] than in the LSA group [mean (SD) = 6.13 (3.43); range: 0 – 11], *t* (43) = −8.62, *p* < .001.

#### Stimuli and Procedure

Pictures of 7 models expressing fearful and neutral expressions were selected from the Karolinska Directed Emotional Faces database (KDEF; (Lundqvist, Flykt, & Ohman, 1998). Fearful and neutral pictures of each model were morphed on a continuum of 0% (pure neutral) to 100% (pure fearful) to create graded fearful expressions (Forscher & Li, 2012). Fear expressions at 6 intensity levels—15%, 21%, 27%, 33%, 39%, and 45%—and a neutral expression, which was set at 2% fear to match visual alterations caused by the morphing procedure, were selected (Forscher & Li, 2012). To include a high number of trials to ensure ERP signal quality, we limited the highest level of fear to 45%, which was determined based on systematic piloting to generate reliable, explicit fear detection. Importantly, as this 45% fear level closely resembled a normative fearful expression in real-life social interactions, it was chosen as the fear prototype here. A total of 686 trials (98 trials per morph level) were presented, randomly intermixed in four blocks, to which participants made two-alternative forced choices (Yes/No) to fear presence.

#### EEG data acquisition & analysis

EEG and Electrooculogram (EOG) was recorded from a 96-channel (ActiveTwo; BioSemi) system at a 1024 Hz sampling rate with a 0.1–100 Hz bandpass filter. EEG/EOG signals were then digitally bandpass filtered from 0.1 to 40 Hz, and down-sampled to 256 Hz. Data were then submitted to Fully Automated Statistical Thresholding for EEG Artifact Rejection (FASTER; (Nolan, Whelan, & Reilly, 2010), a plug-in function in EEGLAB (Delorme & Makeig, 2004). As illustrated in Figure 1 and described in the initial paper (Forscher et al., 2016), our combination of psychophysical testing and neurometric analysis identified four key processes in fear perception—fear categorization, detection, valuation, and conscious awareness— which were respectively indexed by the P1 (at Oz, 82-118 ms centered on the peak latency), the P300 (at Pz, 270-370 ms centered on the peak latency), the early subcomponent of the late positive potential (eLPP; at Pz, 400-500 ms), and the late subcomponent of the late positive potential (lLPP; at Pz, 500-600 ms). As described below, we extracted mean amplitudes of these ERPs and derived respective metrics for the four processes.

#### ERP metrics of key cognitive operations in threat perception

Our sigmoid curve fitting of fear response rates confirmed 2% and 45% fear as neutral and fear prototypes (Figure 2A). The fitted sigmoid curve also denotes 21%, 27%, and 33% fear as intermediate levels and 15% and 39% proximal levels to corresponding prototypes. Incorporating these critical levels, we then derived ERP metrics for the four key operations according to their characteristic neurometric functions (Figure 1). (1) Fear categorization was modelled by an upward quadratic function, depicting maximal responses at prototype levels and minimal responses at intermediate boundary levels (Goldstone & Hendrickson, 2010), and accordingly, the categorization metric was computed as [P1 (Levels 2% + 45%)/2 – P1 (Levels 21% + 27% + 33%)/3]. (2) Fear detection was modeled by a sigmoid function with an asymptote above the detection threshold (21% fear, corresponding to ~25% fear response rate), and the detection metric was accordingly computed as [P300 (Levels 21% + 27% + 33% + 39% + 45%)/5 – P300 Level 2%]. (3) Fear valuation (of fear intensity) was modeled by a linear function of fear intensity, and the valuation metric was computed as [eLPP (3*Level 45% + 2*Level 39% + Level 33% - Level 21% − 2*Level 15% − 3*Level 2%)]. (4) Fear conscious awareness was modeled by a sigmoid function with an asymptote at low to intermediate fear levels until a sharp rise near the threshold of conscious perception (Del Cul, Baillet, & Dehaene, 2007).The awareness metric was accordingly computed as [lLPP (Level 45% + Level 39%)/2 −(Level 2% + Level 15% + Level 21% + Level 27% + Level 33%)/5].

**Figure 2:**
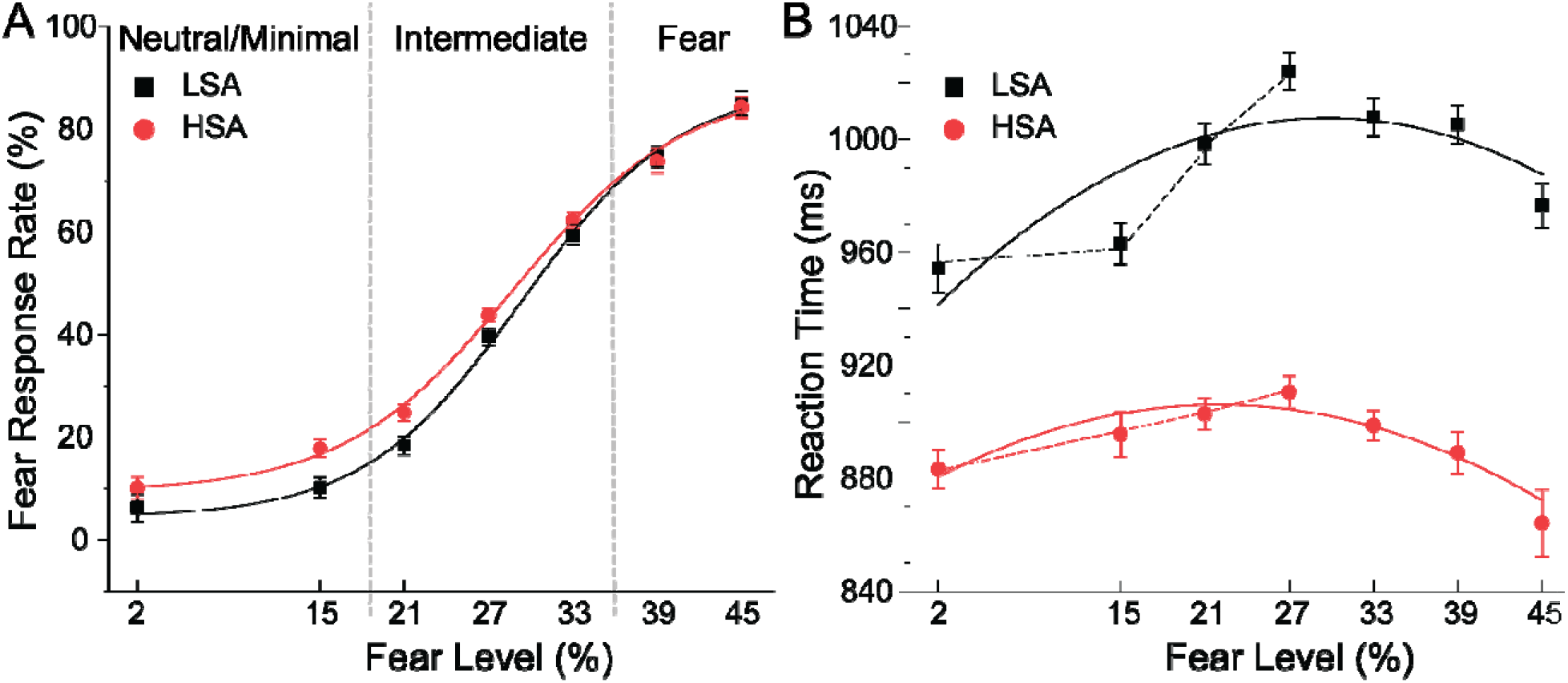
Behavior results. (A) Fear response rate as a function of fear intensity (indexed by fear morph %). The pattern fits very tightly to a sigmoid function for both LSA and HSA groups (black and red solid curve, respectively). (B) Reaction time as a function of fear intensity. The pattern fits a quadratic function for both LSA and HSA groups (black and red solid curve, respectively). The dotted lines illustrate the rather nonlinear vs. linear pattern in the lower half of the continuum for LSA and HSA groups, respectively. Error bars indicate individual-mean-adjusted S.E.M. (i.e., S.E.E.).

#### Statistical analysis

We conducted two-way repeated measures analyses of variance (ANOVAs; with Greenhouse– Geisser corrections) of fear Intensity (fear %) and Group (HSA vs. LSA) on fear response rate and reaction time (RT). Main effects of Intensity were previously reported (Forscher et al., 2016), so here we focused on the simple and interaction effects of Group. We further submitted the ERP metrics for the four key operations, respectively, to between-group *t*-tests. False discovery rate (FDR) was applied to the *p* values to correct for multiple tests. Significant effects were further specified systematically through conventional ANOVAs of Intensity and Group on the ERP amplitude. Finally, we applied curve fitting analyses to the behavioral responses and ERPs (based on the group mean; (Jemel et al., 2003) in OriginPro (OriginLab, Northampton, MA, USA) to determine their fit to the predicted models for each group.

### Results

#### Behavioral Results

##### Fear response rate

An ANOVA of fear Intensity and Group on response rate indicated a main effect of Intensity (as previously reported), but no main effect or interaction of Group (*F*’s < 1.18, *p*’s > .31). Figure 2A revealed a higher fear response rate at the 15% fear level for the HSA than LSA group, which was confirmed by a *post hoc* test (*t* (43) = −2.20, *p* = .03). However, this effect did not survive the FDR correction and will not be discussed further. Response curve fitting indicated strong sigmoid functions of fear intensity for both HSA and LSA groups (*R*^2^ = .99/.99, respectively), clearly separating the prototype and intermediate levels.

##### Fear detection RT

A similar ANOVA on RT revealed a main effect of Intensity (as previously reported). Akin to the binary categorical judgment, response curve fitting confirmed quadratic functions of fear intensity in both HSA and LSA groups (*R*^2^ = .85/.67, respectively), with fastest RT at the prototype levels and slowest RT at the intermediate levels (Figure 2B). In addition, we observed a main effect of Group [*F* (1, 43) = 4.27, *p* = .045, *η*_p_^2^ = .09; faster RT in the HSA than the LSA group] and an Intensity-by-Group interaction [*F* (2.38, 102.19) = 3.25, *p* = .035, *η*_p_^2^ = .07].

To specify this interaction, we performed two ANOVAs of Group and Intensity for the left (2% to 27% fear) and right (27% to 45% fear) halves of the neutral-fear continuum, respectively. A Group-by-Intensity interaction emerged on the left [*F* (2.05, 88.24) = 6.24, *p* = .003, *η*_p_^2^ = .13] but not right half (*p* = .85) of the curve. That is, the groups diverged in their RT profiles from the neutral prototype to the midpoint of the continuum: in the LSA group, RT was fastest at neutral/minimal fear (2% and 15%, which had comparable RT, *p* = .27) and abruptly slowed at the intermediate levels (21% and 27%; pairwise contrasts with neutral/minimal fear levels were all significant, *p*’s < .005), characteristic of categorical emotion perception. By contrast, the HSA group failed to demonstrate categorical perception with a rather flat RT profile; except for a marginal difference between 2% and 15% fear (*p* = .08), RT did not differ between adjacent levels (*p*’s > .27).

### ERP Results

#### Anxiety impaired early fear categorization

The P1 metric of categorization showed an effect of Group [*t*(40) = 2.61, *p* = .013; FDR *p* = .052], reflecting clear categorization in the LSA group [categorization index mean (SD) = .54 (.67); *t*(20) = 3.69, *p* = .001] and impaired categorization in the HSA group [mean (SD) = .09 (.43); *t*(20) = .91, *p* = .37; Figure 3]. Accordingly, curve fitting indicated that the LSA group exhibited a strong quadratic function with maximal P1 amplitudes for the prototypes and minimal P1 amplitudes for intermediate levels (*R*^2^ = .68). By contrast, there was no clear quadratic pattern in the HSA group (*R*^2^ = .19).

**Figure 3:**
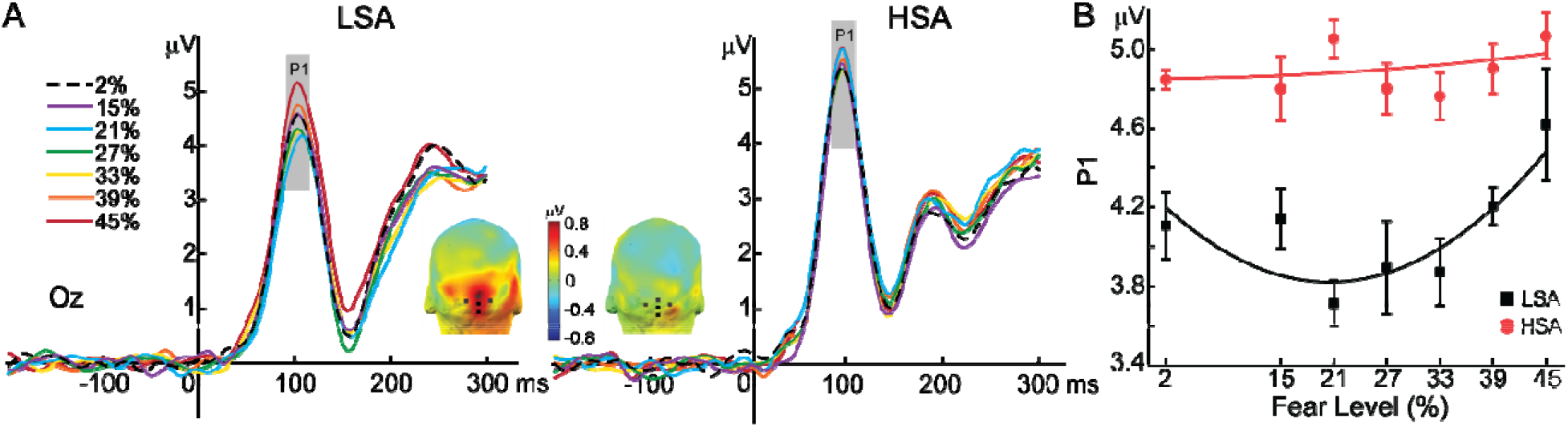
Impaired fear categorization in anxiety. (A) Grand average ERP waveforms at site Oz (marked by 5 black dots in the topographical maps) showing P1 potentials for both LSA and HSA groups. Topographical maps depict the categorization metric [(Levels 2% + 45%) − Levels (21% + 27% + 33%)/3] for each group. (B) P1 amplitude as a function of fear intensity for both LSA and HSA groups. The P1 response pattern conformed to a quadratic fit in the LSA group (black curve) but not the HSA group (red curve). Error bars = S.E.E.

To further specify this effect, we applied a conventional ANOVA (Intensity by Group) on P1 amplitude. We confirmed a significant quadratic Group-by-Intensity interaction effect [*F*(1, 40) = 5.76, *p* = .021, *η*_p_^2^ = .13], which was explained by a strong quadratic effect of Intensity in the LSA group [*F*(1, 20) = 11.40, *p* = .003, *η*_p_^2^ = .36], relative to no effects in the HSA group (*p*’s> .30). Follow-up *t*-tests in the LSA group indicated a smaller P1 for the intermediate levels (pooled across 21%, 27%, and 33%) than the two external levels [the neutral levels (pooled between 2% and 15%): *t*(20) = −2.35, *p* = .029; the fear levels (pooled between 39% and 45%): *t*(20) = −3.33, *p* = .003], while the two external levels showed equivalent P1 amplitudes (*p* = .26).

#### No effect of anxiety on other operations

As illustrated in Figure 4, profiles of the other three ERP components conformed to the predicted response functions for both groups: strong sigmoid fit for P300 and late LPP components and strong linear fit for the early LPP component, *R*^2^ > .87), with the exception of a poor sigmoid fit for P300 in the LSA group (*R*^2^ = .20; Figure 4B). Nonetheless, there was no group effect on these ERP metrics (fear detection, valuation, or conscious awareness), *t*’s < 1.22, *p*’s > .23.

**Figure 4:**
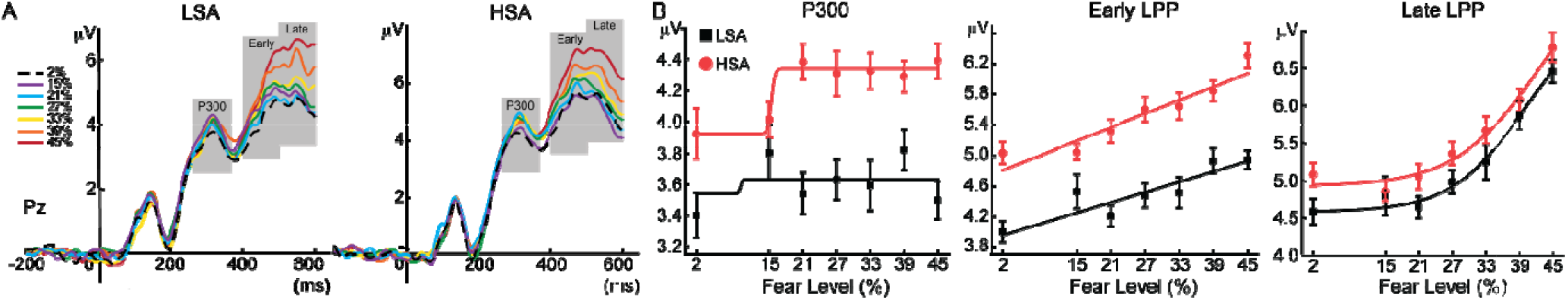
Fear detection, valuation, and conscious awareness. (A) Grand average ERP waveforms at site Pz showing P300, early LPP, and late LPP components for LSA (Left) and HSA (Right) groups. (B) ERP metrics for fear detection (P300), valuation (early LPP), and conscious awareness (late LPP) largely conformed to their characteristic response curves, with the exception of P300 in the HSA group. Note, although raw amplitudes for these ERPs appeared to be larger in the HSA (vs. LSA) group, these group differences failed to reach statistical significance, *F*’s < 2.14, *p*’s > .15. Error bars = S.E.E.

## Discussion

Social anxiety is associated with biased social perception, especially of ambiguous signals. By combining psychophysical and neurometric analyses among individuals with high or low trait social anxiety, we identified a dispositional impairment in categorical perception of threat that could contribute to such biases in social anxiety. Clear divergence between high- and low-social anxiety groups emerged in the RT profile along the neutral-fear continuum—anxious individuals failed to demonstrate a profile characteristic of categorical perception that was nonetheless manifested by non-anxious individuals. Confirming the bias in social anxiety that is most salient for ambiguous threat, the disparity in the RT profile was especially pronounced from neutral to mild/moderate fear intensities. In parallel, the neural (P1) metric (but not later operations) isolated an early fear categorization process at 100 ms in the non-anxious (but not anxious) individuals. Therefore, behavioral and neural evidence converged to demonstrate impaired categorical perception of threat in trait social anxiety, implicating an early, automatic perceptual mechanism in the pathology of social anxiety.

Akin to hypervigilance in anxiety, the HSA group exhibited overall faster responses than the LSA group. On top of this general trend, RT profiles, across the 7 intensity levels along the neutral-fear continuum, conformed to categorical threat perception in the LSA group but aligned with dimensional threat perception in the HSA group. Specifically, LSA individuals showed greater ease (faster RT) in response to neutral/minimal fear levels (2%/15%) in contrast to abruptly increased difficulty (much slower RT) to intermediate levels (21-33%), suggesting a clear distinction between no threat versus ambiguous cues (or potential threat). However, HSA individuals showed monotonous and small (non-significant) RT increases across these levels, failing to draw such a boundary. It is conceivable that the breakdown of such an important categorical boundary and the consequent impairment in ambiguity resolution would subject anxious individuals to a persistent presence of potential threat in their social environment, fueling and perpetuating their symptoms.

Our recent parametric modeling of ERP responses along the fear continuum has captured four key cognitive operations (Forscher et al., 2016). Here, leveraging these neurometric functions, we derived ERP metrics for the strength of these operations and demonstrated that the operation of threat categorization was impaired in the anxious group. While the LSA group exhibited a quadratic function in P1 responses across the fear levels, the HSA group displayed a rather flat, linear profile. Conforming to the neural pattern of stimulus categorization (i.e., pronounced responses to the prototypes vs. suppressed response to boundary stimuli; (Goldstone & Hendrickson, 2010), the LSA group responded equally and strongly to the external/prototype stimuli (neutral/15% vs. 39%/45%) but weakly to intermediate stimuli (21%-33% fear). Essentially, it was this response suppression to intermediate fear stimuli in the LSA group and the lack thereof in the HSA group that set the two groups apart.

The P1 component, originating in the extrastriate cortex around 100 ms post-stimulus, is a reliable index of early visual processing (Gomez Gonzalez, Clark, Fan, Luck, & Hillyard, 1994; Mangun, Hillyard, & Luck, 1993). Ample research has demonstrated differential P1 response to threat versus non-threat stimuli, with its intracranial sources localized to early visual cortices in the occipital lobe (cf. (W. Li, 2019; Miskovic & Keil, 2012; Vuilleumier & Pourtois, 2007). This P1 differentiation of threat (vs. non-threat) could be especially pronounced in anxious individuals, which is often assumed to reflect arousal or attentional bias to threat (Eimer & Holmes, 2007; Forscher & Li, 2012; Krusemark & Li, 2011; W. Li, Zinbarg, et al., 2008; W. Li, Zinbarg, & Paller, 2007; Rossignol, Campanella, Bissot, & Philippot, 2013; Weinberg & Hajcak, 2011). However, new evidence of differential P1 to subtypes of threat stimuli (e.g. disgust vs. fear/anger) and findings of reduced P1 to disgust/disliked (relative to neutral/liked) stimuli are incompatible with this assumption (Krusemark & Li, 2011, 2013; Liu, Zhang, & Luo, 2015; Pizzagalli, Regard, & Lehmann, 1999; You & Li, 2016), implicating a more complex process (beyond simple arousal or attention modulation) in this early visual processing of threat.

In standard object perception (purportedly independent of emotion-related arousal or attention), differential P1 responses are observed for different categories (e.g., indoor vs. outdoor scenes) and are thought to reflect stimulus categorization (Thorpe, 2009; Thorpe et al., 1996). Here, the strong quadratic function of P1 responses (with equivalent response to neutral and clearly fearful expressions) in non-anxious individuals is more aligned with the idea of categorization (of fear or non-fear faces) than mere arousal/attention-related response modulation. In addition, this categorization process differs from fear detection, which immediately follows as reflected in the P300 component, in its binary (vs. singular) classification of threat versus non-threat/safety. Reflecting an initial, coarse threat classification, it is also distinct from later processes associated with deliberate and nuanced categorization (i.e.., valuation of threat level and conscious awareness of threat as indexed by the LPP components).

This early binary (threat or safety) process coincides with computational models of “saliency maps” (Z. Li, 2002) and neuroscience models of salience detection (Menon & Uddin, 2010; Seeley et al., 2007), implicating fast isolation of biologically meaningful signals from noise (stimuli to be dismissed) during early sensory processing, prompting efficient responses to salient events. In addition, threat encoding in the sensory cortex could further furnish this early categorization process with direct cortical input (W. Li, 2014; W. Li, Howard, Parrish, & Gottfried, 2008; McTeague, Gruss, & Keil, 2015). In sum, we speculate that coarse emotion categorization could be triggered by automatic, bottom-up sensory input (Brosch, Pourtois, & Sander, 2010; Young et al., 1997) and culminate at an early stage by integrating sensory emotion encoding with basic sensory processes (e.g. basic feature processing, template matching; (Krusemark & Li, 2011, 2013; You & Li, 2016).

Nonetheless, this ecologically advantageous process is impaired in socially anxious individuals, largely due to diminished response suppression at intermediate fear levels. This impairment in early sensory suppression of ambiguous, boundary cues is consistent with the notion of sensory disinhibition in anxiety (Clancy, Ding, Bernat, Schmidt, & Li, 2017; W. Li, 2019). That is, while anxiety is characterized by heightened response to threat, it has also been associated with broad (*threat-neutral*) enhancement (or disinhibition) of early sensory processing. Patients with post-traumatic stress disorder (PTSD) exhibit attenuated P50 suppression to double-click stimuli (reflecting sensory gating impairment) and exaggerated auditory and visual ERPs to simple, neutral stimuli (e.g., a tone); reflecting sensory cortical hyperactivity (Javanbakht, Liberzon, Amirsadri, Gjini, & Boutros, 2011; Lewine et al., 2002; Morgan & Grillon, 1999; Neylan et al., 1999; Skinner et al., 1999). Spider phobics demonstrate comparable exaggeration of visual ERPs (P1 and C1) to images of spiders and unrelated objects (Michalowski et al., 2009; Michalowski, Pane-Farre, Low, & Hamm, 2015; Michalowski, Weymar, & Hamm, 2014). Of particular relevance here, besides aforementioned evidence of *specific* P1 enhancement to threatening faces, there is almost equally strong evidence of *generic* P1 enhancement to faces, regardless of facial expressions, in social anxiety (Helfinstein, White, Bar-Haim, & Fox, 2008; Kolassa et al., 2009; Kolassa, Kolassa, Musial, & Miltner, 2007; Kolassa & Miltner, 2006; Muhlberger et al., 2009; Peschard, Philippot, Joassin, & Rossignol, 2013; Rossignol, Campanella, et al., 2012; Rossignol, Philippot, Bissot, Rigoulot, & Campanella, 2012; Wieser & Moscovitch, 2015). Therefore, anxiety could be associated with sensory disinhibition, which would compromise early stimulus categorization, resulting in biased threat perception. Notably, our parametric delineation of anxiety modulation of P1 responses may provide useful insights into the mixed findings in the literature, implicating variability in threat intensity as a source for the sometimes specific and sometimes generic effects of anxiety.

In conclusion, by modeling fear processing across a neutral-fear continuum, we identified impaired threat categorization in association with trait social anxiety, which arises from disinhibited early sensory response to ambiguous cues. This lack of early sensory inhibition towards dismissible signals could represent a failure to filter (or “gate”) out innocuous sensory input from entering downstream processing, triggering excessive threat responses (Clancy et al., 2017; W. Li, 2019). In the absence of significant effects of trait anxiety on later operations, impaired early categorization of threat may reflect a dispositional cognitive vulnerability, predisposing an individual to further cognitive aberrations and, eventually, clinical symptoms of anxiety.

## Acknowledgements

This research was supported by the National Institute of Mental Health (R01MH093413 to W.L.).

